# Modules of co-occurrence in the cyanobacterial pan-genome

**DOI:** 10.1101/137398

**Authors:** Christian Beck, Henning Knoop, Ralf Steuer

## Abstract

The increasing availability of fully sequenced cyanobacterial genomes opens unprecedented opportunities to investigate the manifold adaptations and functional relationships that determine the genetic content of individual bacterial species. Here, we use comparative genome analysis to investigate the cyanobacterial pan-genome based on 77 strains whose complete genome sequence is available. Our focus is the co-occurrence of likely ortholog genes, denoted as CLOGs. We conjecture that co-occurrence CLOGs is indicative of functional relationships between the respective genes. Going beyond the analysis of pair-wise co-occurrences, we introduce a novel network approach to identify modules of co-occurring ortholog genes. Our results demonstrate that these modules exhibit a high degree of functional coherence and reveal known as well as previously unknown functional relationships. We argue that the high functional coherence observed for the extracted modules is a consequence of the similar-yet-diverse nature of the cyanobacterial phylum. We provide a simple toolbox that facilitates further analysis of our results with respect to specific cyanobacterial genes of interest.

## INTRODUCTION

Cyanobacteria are photosynthetic prokaryotes of global importance and recently gained renewed interest as a resource for natural products (Abed et al. 2009; Calteau et al. 2014) and as potential host organisms for the renewable synthesis of bulk chemicals and including biofuels (Gupta et al. 2013; Cassier-Chauvat et al. 2014; Savakis and Hellingwerf 2015). Cyanobacteria exhibit diverse morphologies and are known to inhabit diverse environments, including lakes, oceans, arctic rocks, desert crusts, hot springs, and rice fields. In addition to using oxygenic photosynthesis as a primary source of energy and reducing power, many cyanobacteria are capable to assimilate atmospheric nitrogen, making cyanobacteria key players in the global nitrogen cycle. Despite their ecological and biotechnological importance, however, many aspects of cyanobacterial diversity are still insufficiently understood. In this respect, the increasing availability of fully sequenced cyanobacterial genomes (Shih et al. 2013; Fujisawa et al. 2017) opens unprecedented opportunities to delineate cyanobacterial diversity and the physiological adaptations from a genomic perspective.

Comparative genome analysis, in particular the analysis of the bacterial pan-genome, is established for more a decade (Haft 2015; Vernikos et al. 2015). Early applications include a comparison of 8 Streptococcus genomes (Tettelin et al. 2005), followed by an analysis of 17 *Escherichia coli* genomes (*Rasko et al. 2008*), and similar studies for *Legionella pneumophila* (D’Auria et al. 2010), *Haemophilus influenzae* (Hogg et al. 2007), and twelve closely related strains of the cyanobacterial genus *Prochlorococcus* (Kettler et al. 2007). Later studies considered an increasing number of strains (Donati et al. 2010; Beck et al. 2012), as well as an analysis of niche-specific differences (Simm et al. 2015). Several computational approaches and toolboxes for comparative genome analysis and the identification of ortholog genes have been described in the literature (Fouts et al. 2012; Vallenet et al. 2013; Waterhouse et al. 2013; Benedict et al. 2014; Galperin et al. 2015).

In this work, we report a comparative analysis of 77 fully sequenced and assembled cyanobacterial genomes. Beyond the analysis of the pan- and core-genome, we are particularly interested in the co-occurrence of Clusters of Likely Ortholog Genes (CLOGs). Specifically, we hypothesize that genes with related functions also co-occur within genomes. A well-known example of such co-occurrences are operons, sets of genes that are under control of a single promotor and act as a functional unit (Jacob et al. 1960). Operons play a significant role in the parallel inheritance of physiologically coupled genes through horizontal gene transfer (Koonin et al. 2001; Pål et al. 2005). More general, however, sets of genes that constitute a functional unit must not necessarily be co-localized or be under the control of a single promotor. Nonetheless, a functional dependency typically still requires co-occurrence within the same genome. While pair-wise co-occurrences can be straightforwardly detected, the identification of larger functional units, denoted as modules of co-occurring genes, involves the analysis of the community structure of large networks -- a computationally nontrivial task. In the following, we demonstrate that an analysis of co-occurrences indeed provides hypotheses about functional relationships between genes and augments established analysis based on co-localization and other criteria.

The paper is organized as follows. In the first two sections, we briefly recapitulate several key properties of the cyanobacterial pan- and core genome. In the third section, we then identify co-occurring CLOGs within the cyanobacterial pan-genome. In the following section, we then go beyond pair-wise analysis and introduce a weighted co-occurrence network that is used to identify groups (modules) of co-occurring CLOGs. In the fifth section, we demonstrate that co-occurrence does not imply co-localization. The final sections are then devoted to analyze selected examples of modules of co-occurring CLOGs. We demonstrate that these modules indeed correspond to functional relationships and provide novel hypotheses for gene function. To facilitate further analysis, the work is supplemented with the software toolbox “CyanoCLOG SimilarityViewer” to allow for the exploration of co-occurrences beyond the selected examples discussed here.

## RESULTS

### The cyanobacterial pan-genome revisited

Starting point of our analysis are 77 sequenced cyanobacteria sourced from the NCBI GenBank database. To avoid bias due to incomplete genomes, only completely assembled chromosomes, together with their associated plasmids were selected (132 plasmids total). For later reference, the strain *Escherichia coli O111*:*H* (denoted as E. coli in the following) was included within the analysis.

Orthology of all identified genes was determined based on an all-against-all BLASTp search as described previously (Beck et al. 2012). Gene pairs with a high BLAST score and bidirectional hit rate (BHR) were grouped together and subsequently clustered based on their mutual BLAST score into cluster of likely orthologous genes, denoted as CLOGs. See METHODS for computational details. Due to their unique properties, the cyanobacterium UCYN-A, an endosymbiont with a highly reduced genome (Zehr et al. 2008), and E. coli were not part of the initial core- and pan-genome analysis. Their CLOGs were kept for later analysis. We distinguish between core CLOGs, present in all remaining 76 strains, shared CLOGs, present in one or more but not in all strains, and unique CLOGs, identified only in a single strain. Overall, we identified a total of 58740 CLOGs consisting of 621 core CLOGs, 20005 shared CLOGs, and 38114 unique CLOGs. Strains with larger genomes tend to be associated with more shared CLOGs. In contrast, the number of unique CLOGs associated with a single strain is dependent also on the phylogenetic distance to its nearest neighbors -- and is therefore biased by the coverage of the cyanobacterial phylum, rather than being an intrinsic property of a strain. The number of strains associated with each CLOG is shown in Figure 1A.

The overall properties of the pan- and core-genome are in good agreement with previous studies, typically using a smaller number of strains (Beck et al. 2012; Simm et al. 2015). Core CLOGs constitute between 7.4% (Acaryochloris marina MBIC11017) and 33.5% (Prochlorococcus marinus str. MIT 9211) of all CLOGs in a given genome. Evaluating the pan-genome for a subset of strain allows us to extrapolate the expected increase in the size of the pan-genome for newly sequenced genome. The data follows Heap’s law (Figure 1B), indicating approximately 450 genes with sub-threshold similarity to any known protein for each newly sequenced genome (Tettelin et al. 2008). These numbers are in good agreement with the 21,107 novel sub-threshold genes identified by recent de-novo sequencing of 54 cyanobacterial strains (Shih et al. 2013). We note that extrapolation of the core genome (Figure 1C) should be interpreted with caution due to the inherent statistical caveats when estimating rare events from limited data (Kislyuk et al. 2011).

**Figure 1:**
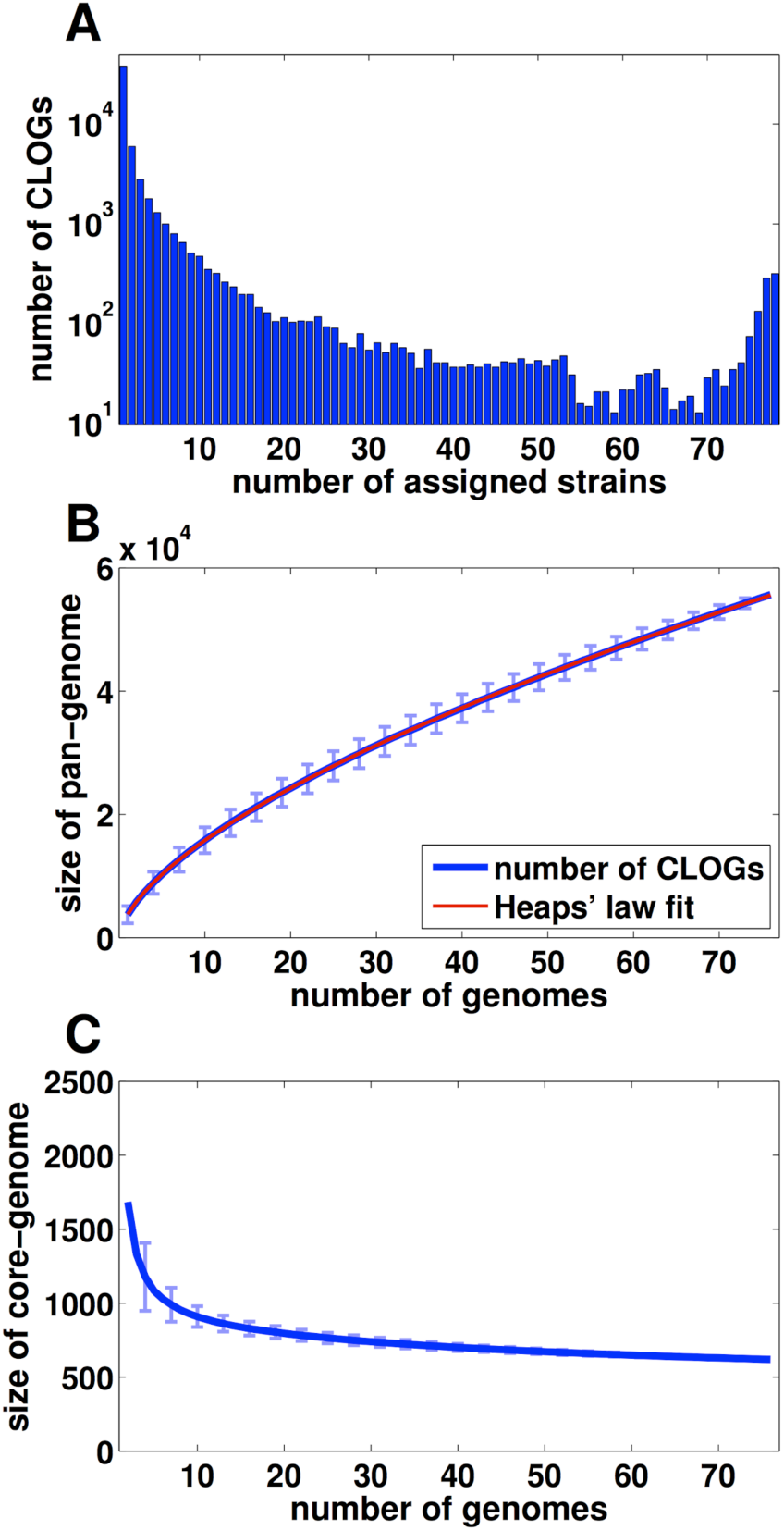
The cyanobacterial pan- and core-genome. (A) Distribution of CLOGs as a function of the number of assigned strains. (B) Size of the pan-genome estimated for an increasing number of strains. The blue line indicates the mean size of the pan-genome, error bars indicate the standard deviation of 10^4^ randomly sampled subsets of strains. The red line shows a least squares fit of the power law p ∼ N^g^ (Heaps’ law), with p denoting the size of pan-genome and N the number of genomes. The estimated exponent g = 0.62 indicates an open pan-genome. (C) Size of the cyanobacterial core-genome estimated for an increasing number of strains. The blue line indicates the mean size of the core-genome whereas error bars indicate the standard deviation of 10^4^ randomly sampled subsets of strains. The estimates of pan- and core-genome do not include genomes of E. coli and *Cyanobacterium* UCYN-A.

### CLOG annotation and the cyanobacterial pan-metabolism

CLOGs can be assigned to functional categories based on the annotation of their constituent genes. As expected, annotation coverage in core CLOGs is high, with 589 of 621 (95%) of core CLOGs containing at least one gene with functional annotation (as sourced from GenBank, including unspecific annotation, such as *membrane protein* but excluding annotations such as *hypothetical* and *conserved hypothetical*). Consistent with earlier studies (Mulkidjanian et al. 2006; Beck et al. 2012; Simm et al. 2015), functional annotation of core CLOGs is enriched in categories related to cellular metabolism, transcription, translation, and DNA replication. As compared to core CLOGs, annotation coverage for shared and unique CLOGs is significantly lower with 44% (8853 of 20005) and 82% (31132 of 38114), respectively, annotated as *hypothetical*, *predicted* or *unknown*. As observed previously (Beck et al. 2012), annotation is often unspecific or varying in the exact wording, for example “*photosystem II DII subunit”* and “*photosystem II protein D2”*. Yet we observe only few instances of conflicting annotations for two or more genes in one CLOG. While automated comparison of conflicting annotation is not straightforward, manual inspection of 1000 CLOGs with at least two genes revealed putative inconsistencies for only 2.5% of the CLOGs. That is, in less than three percent of cases with at least two semantically different annotations, the annotations could not be identified as coinciding at a first glance.

We are specifically interested in the cyanobacterial pan-metabolism. To this end, the constituent genes of each CLOG were matched against the KEGG (Kyoto Encyclopedia of Genes and Genomes) database (Kanehisa et al. 2008) to identify, which CLOGs are associated to EC numbers. We obtained a total of 2361 metabolism-related CLOGs associated with a total of 2301 metabolic reactions. We note that enzymes (and hence CLOGs) may catalyze multiple reactions and multiple enzymes (and hence CLOGs) may catalyze the same reaction. Core CLOGs are highly enriched in metabolic function, with 322 of 621 (51.9%) associated with one or more specific reaction. The ratio is significantly lower for shared and unique CLOGs, with only 1664 (8.3%) shared CLOGs and 408 unique CLOGs (1.1%) associated to one or more specific reactions. Nonetheless, due to the higher number of shared CLOGs, metabolic functionality is primarily encoded in the shared genome. Of the 2301 unique reactions that constitute the cyanobacterial pan-metabolism, 1839 reactions are associated with at least one shared CLOG.

### Co-occurring CLOGs indicate functional relationships

We seek to identify putative functional relationships between CLOGs and hypothesize that co-occurring CLOGs are indicative of a possible functional relationship. We first performed a right-tailed Fisher’s exact test to identify pairs of CLOGs who preferentially co-occur within the same strain (see METHODS). Using the multiple test correction method for non-independent tests by Benjamini and Yekutieli (Benjamini and Yekutieli 2001), we obtained a critical p-value of 1.43*10^-6^ for an accepted false discovery rate (FDR) of 0.01. In consequence, we identified 581,741 out of more than 1.7*10^9^ possible pairs of CLOGs whose occurrences are significantly correlated. We note that, by definition, co-occurrence only relates to shared CLOGs. Core CLOGs are not considered in the analysis. The full list of co-occurring CLOGs is provided in Supplement Table 1.

Manual inspection of co-occurring CLOGs indeed indicates functional relationships. For example, among the pairs with the lowest p-value are the two subunits of cytochrome bd plastoquinol oxidase (CLOGs 11458 and 11459), typically forming an operon. Another example is the co-occurrence between subunits of the hydrogenase maturation protein Hyp, specifically the co-occurence of HypA (CLOG 10002) and HypE (CLOG 12374), as well as of HypE and HypF (CLOG 11744). Importantly, these subunits are not in close proximity on the genome in about half of all strains (16 of 39). Likewise, genes that encode a tocopherol cyclase (EC 5.5.1.24, CLOG 9703) and a homogentisate phytyltransferase (EC 2.5.1.115, CLOG 10825) co-occur. Both CLOGs also co-occur with a CLOG encoding a 4-hydroxyphenylpyruvate dioxygenase (EC 1.13.11.27, CLOG 10837). The genes of this triplet are essential for the biosynthesis of Vitamin E but are not in close genomic proximity on any strain. In the following, we therefore address two aspects of co-occurring CLOGs. Firstly, functional relationships go beyond pairs and may involve groups of CLOGs. Secondly, the genes of co-occurring CLOGs are not necessarily always in close proximity on the genome. Since co-localization of genes on genomes of different species is already used for functional analysis (Winter et al. 2016), the latter fact indicates that co-occurrence provides additional information that augments co-localization as an indicator of a functional relationship.

We note that, in addition to co-occurrence, also anti-occurrence can be studied. Mutually exclusive pairs of CLOGs, however, are typically associated with specific phylogenetic clades, such as an exclusive association to either alpha-cyanobacteria or beta-cyanobacteria. In the following, we therefore focus on co-occurrence only. A brief analysis of anti-correlated pairs of CLOGs is provided in the Supplementary Text 1.

### Network analysis of co-occurring CLOGs

Pair-wise co-occurrences are not sufficient to fully reveal the underlying structure of functionally related CLOGs. Therefore we seek to identify groups of CLOGs, denoted as *modules*, that co-occur across different genomes. To this end, we consider CLOGs as nodes in a network, such that two CLOGs are connected by a (weighted) link if their co-occurrence is statistically significant. We utilize the weight function w(a,b) between two co-occurring CLOGs a and b,

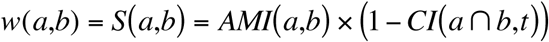

where *AMI*(*a*,*b*) denotes the adjusted mutual information between the occurrences of CLOGs a and *b*, and *CI*(*a ⋂ b*,*t*) denotes the consistency index with respect to the phylogenetic tree. The adjusted mutual information AMI provides a measure for co-occurrence between zero and unity that is corrected for CLOG size (Vinh et al. 2010). The consistency index CI measures the phylogenetic distance of the participating genomes, such that links between CLOGs that co-occur in phylogenetically closely related genomes (CI ∼ 1) have less weight, compared to links between CLOGS that co-occur in phylogenetically distant genomes (CI ∼ 0). The CI adjusts for the fact that co-occurrence is biased by phylogenetic proximity. See METHODS for details.

Based on the resulting weighted co-occurrence network, modules of co-occurring CLOGs were identified using the algorithm of Blondel et al. (Blondel et al. 2008). The algorithm is based on heuristic modularity optimization, parameter-free and reasonably fast. We note that module identification is computationally hard and precise (non-heuristic) formulations are computationally intractable for large networks. Input to the algorithm is the weighted co-occurrence network using a value of w=0.65 as specific cutoff for minimal weight of edges. The results are highly robust with respect to different choices of the cutoff. The workflow is depicted in Figure 2.

The algorithm of Blondel results in 563 modules comprising a total of 1930 CLOGs. Most modules (542 of 563) are of size ten or less, 93 modules consist of three, 371 modules consist of only two CLOGs, respectively. All identified modules and their constituent CLOGs are listed in Supplement Table 2. We note that, despite the correction for phylogenic proximity, large modules typically reflect different subgroups of cyanobacteria. For example, the largest module consists of 48 CLOGs with seemingly unrelated functional annotation, who (co-)occur in most ß-cyanobacteria, with the exception of both *Gloeobacter* strains. Similar, the second largest module (41 CLOGs, of which 25 have no annotated function) is mostly associated with alpha-cyanobacteria, excluding all but two Prochlorococcus strains. Smaller modules, however, are typically not associated to particular clades or subtrees and indicate functional relationship between CLOGs.

**Figure 2:**
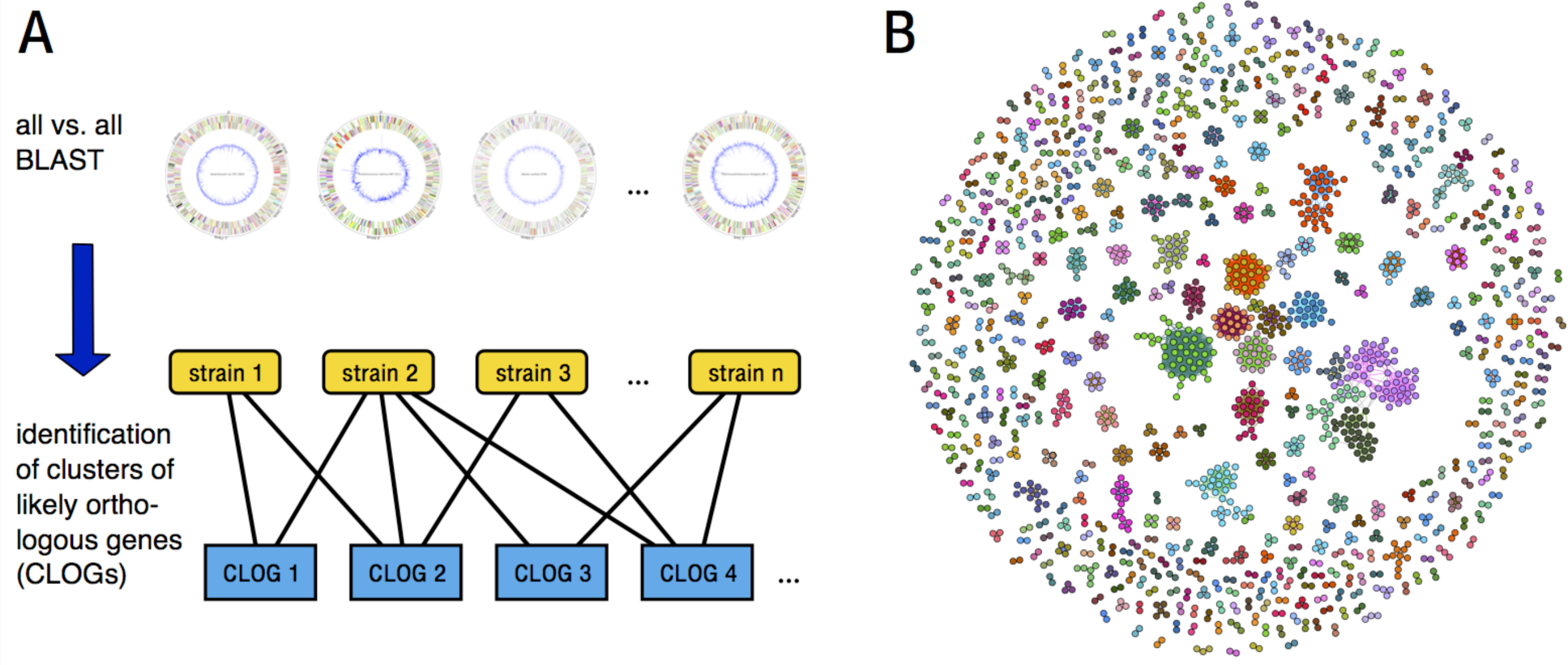
Network analysis of co-occurring CLOGs. (A) Orthologous genes are identified using an exhaustive all versus all BLAST search and subsequently grouped to Cluster of Likely Orthologous Genes (CLOGs). CLOGs can be classified into three sets: core CLOGs (present in all strains), shared CLOGs (present in some strains) and unique CLOGs (present in a single strain). Two shared CLOGs, a and b, may co-occur in different strains, such that CLOG a is annotated within a strain if and only if CLOG b is also annotated within this strain. (B) A network view on co-occurring CLOGs. CLOGs are interpreted as nodes and connected by a weighted edge if both CLOGs co-occur (using a threshold of w > 0.65). Network analysis reveals the community structure of the co-occurrence network and allows to identify modules of co-occurring CLOGs. Circular maps in (A) were constructed using the CiVi tool (Overmars et al. 2015).

### Co-occurrence and co-localization

Prior to a detailed analysis of putative functional relationships between CLOGs, we evaluate to what extend the modules of co-occurring CLOGs reflect co-localization in the genome. In particular, we seek to ascertain that modules do not merely recapitulate the known operon structure of functionally related CLOGs. To this end, we tested the modules of co-occurring CLOGs for operon-like structures. For each module an average adjacency score (aAS) was calculated. For each strain, the strain-specific adjacency score of a module is AS = 1 if all corresponding genes within the module are located in close proximity in the respective genome (defined as being separated by less than ten open reading frames from each other), and AS = 0 if no two genes within a module are located in close proximity to each other. The average adjacency score (aAS) of a module is then given as the average of the AS of all constituent strains. See METHODS for details. The distribution of the aAS for each module is shown in Figure 3A. We observe a dichotomy between modules whose constituent CLOGs (and hence genes) are co-localized in all genomes (aAS ∼ 1) and modules whose genes are not co-localized (a AS ∼ 0). The strict dichotomy is partly explained by the fact that a large number of modules (371 of 563) consist only of two CLOGs, hence the respective AS can be either zero or one. Interestingly, the quality of co-occurrences within a module, measured by the average weight function, shows only a weak correlation with genomic adjacency (Figure 3B). Modules with a low average weight function between its CLOGs may exhibit a similar range of adjacency scores as modules with high average weight function.

**Figure 3:**
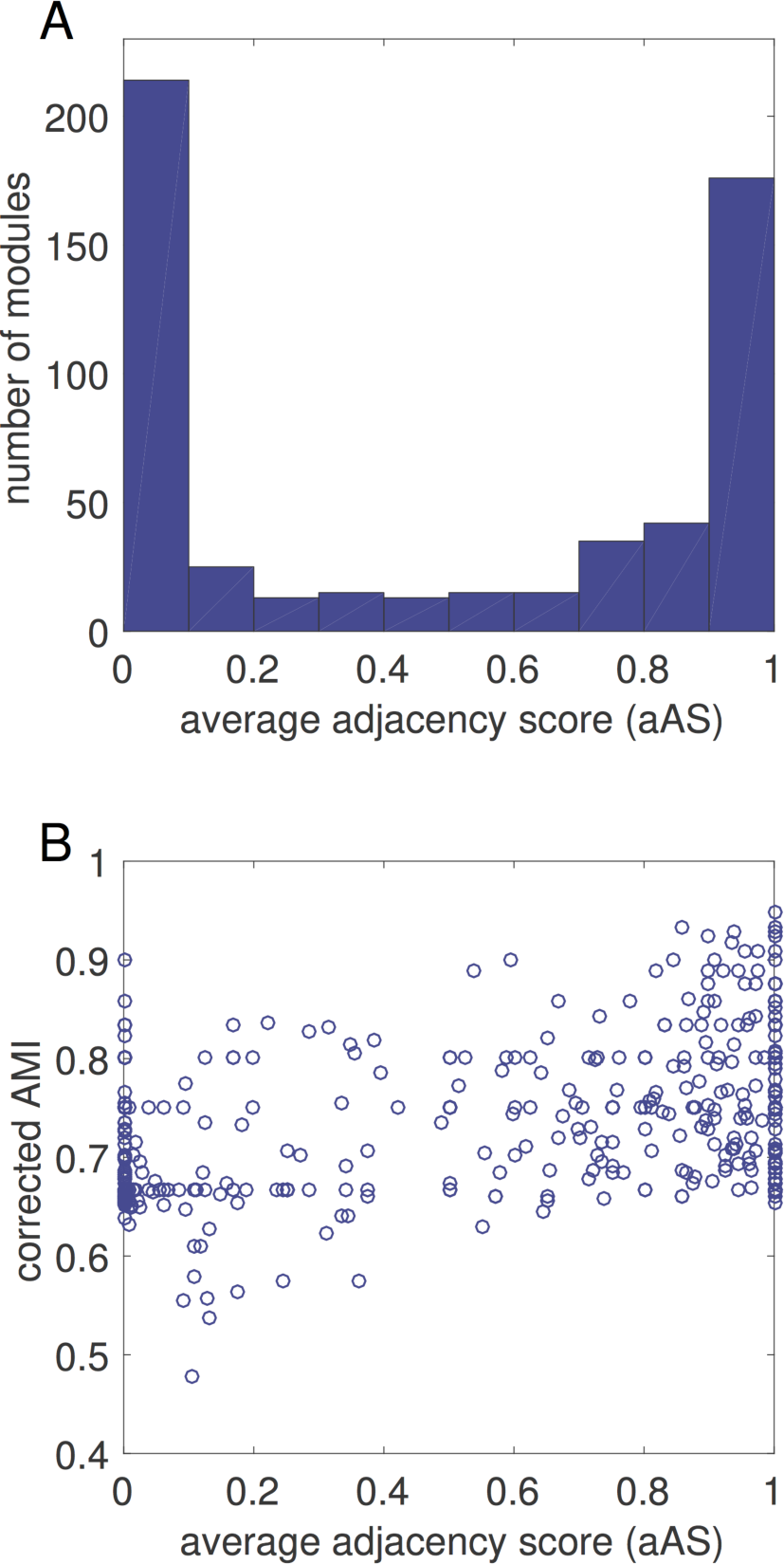
Genomic proximity of co-occurring CLOGs. The adjacency score represents the proximity of CLOGs grouped in modules on the genomic level. The histogram in (**A**) shows a clear dichotomy of the average adjacency score. The scatter plot in (**B**) reveals no correlation between the average adjacency score of a module and the quality of the co-occurrence, represented by the corrected adjusted mutual information.

In general, modules with high adjacency score often correspond to known operons. For example, module 9 (aAS=0.87) contains 21 CLOGs related to the formation of nitrogenase (EC 1.18.6.1), whose genes are arranged in one to three operon-like groups in the genome of most strains. The operon structure of nitrogenase-related genes was previously described by Mulligan and Haselkorn (Mulligan and Haselkorn 1989). But modules with lower AS may also consist of CLOGs that functionally closely related. For example module 293 (aAS = 0) consists of two CLOGs that are annotated as a substrate-binding and membrane subunit of a carbohydrate ABC transporter. The co-occuring subunits are not in close proximity in any cyanobacterial genome.

We note that the genomic proximity of genes is rather conserved in general, with 317 of 563 modules having the same AS in all associated strains, but it can also vary drastically between strains. For example module 52 (aAS=0.34) consists of six genes for nitrate reductase (EC 1.7.7.2) and five proteins associated with its assembly. Despite their close functional relationship, the corresponding genes are organized in operons-like structures in only 13 strains (mostly *Synechococcus*) but are spread across the genomes of 21 other strains. Variations of the genomic adjacency between strains are not straightforward. Neither smaller, streamlined genomes, nor strains with genes organized in multiple plasmids feature a generally difference in the number of operon-like structures (see Supplement Text 1). Overall we conclude, that analysis of the genomic neighborhood is not sufficient to determine candidates for functionally related genes, and that modules of co-occurring genes do not merely recapitulate known co-localization.

### Modules of co-occurring CLOGs indicate functional relationships

To evaluate to what extent modules of co-occurrence provide novel hypotheses for functional relationships between CLOGs, we exemplarily discuss 20 identified modules in the following. The relationship between the selected modules and their constituent CLOGs are depicted in Figure 4.

**Figure 4:**
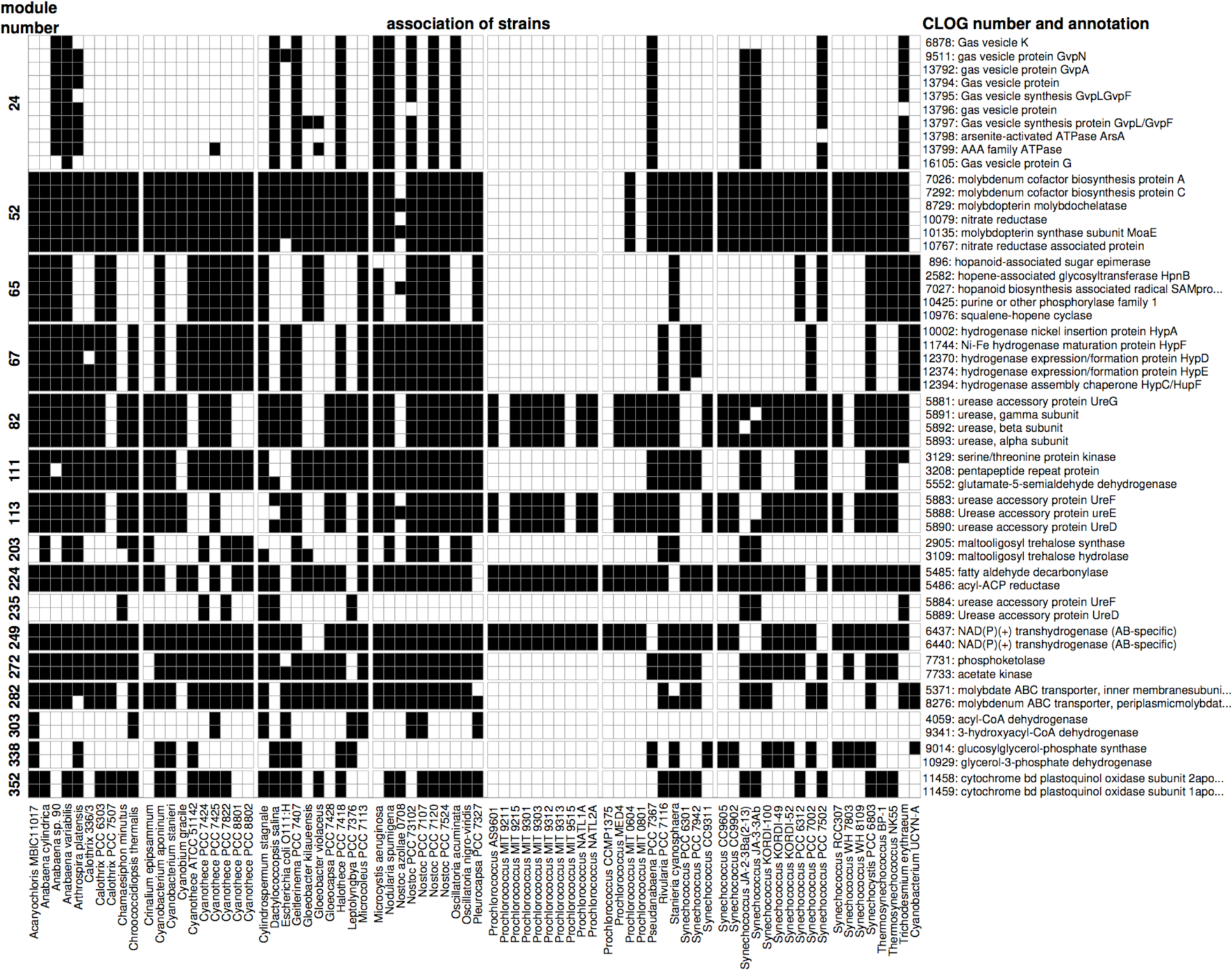
Selected modules and their associated strains. A black box indicates if a CLOG (y-axis) is associated with a specific strain (x-axis). The first column indicates the module number, the last column indicates the primary annotation of the respective CLOG. Shown is an excerpt of modules of co-occurring CLOGs.

The most straightforward instances of functional relationships between CLOGs are subunits of heteromultimeric proteins that co-occur across diverse genomes. For example, **module 249** (2 CLOGs, aAS=0.99) consists of two CLOGs coding for the alpha and beta subunit of a NAD(P)+ transhydrogenase (EC 1.6.1.2) and **module 352** (2 CLOGs, aAS=1) consists of the subunits I and II of cytochrome bd quinol oxidase (EC 1.10.3.14) (Howitt and Vermaas 1998). **Module 67** (5 CLOGs, aAS=0.73) consists of the NiFe-type hydrogenase maturation protein subunits HypA, HypC, HypD, HypE and HybF (Casalot and Rousset 2001). Interestingly, the sixth subunit HypB (CLOG 5882) is not present in this module. The subunit can often be found in multiple copies and is also present in cyanobacteria that do not harbor the other 5 subunits (e.g. *Leptolyngbya* PCC 7376, *Synechococcus* CC 9605, among others), suggesting a possible second function. **Module 82** (4 CLOGs, aAS=0.68) consists of the alpha, beta, and gamma subunits of urease (EC 3.5.1.5) as well as the urease accessory protein UreG. Urease catalyzes the hydrolysis of urea into carbon dioxide and ammonia as a source of nitrogen. The protein complex assembly is assisted by the four chaperons UreD, UreE, UreF, and UreG (Musiani et al. 2004; Fong et al. 2013). For most strains, the accessory proteins UreD, UreE, and UreF are grouped in **Module 113** (3 CLOGS, aAS=0.68). The remaining cyanobacteria that possess urease (e.g. Cyanothece PCC 7424, Leptolyngbya PCC 7376, Trichodesmium IMS101) have a modified UreD and UreF (**Module 235,** 2 CLOGS, aAS=0.78) but lag the UreE chaperon.

A second class of functional relationships is modules whose constituent CLOGs encode transporters. In cyanobacteria and other gram-negative bacteria, ABC (ATP binding cassette) transporters are usually comprised of 3 different molecular components: an ATP-binding/ hydrolyzing protein (NBD - nucleotide binding domain), one or two transmembrane proteins (TMD) building a homo- or heterodimeric structure, and a soluble, secreted substrate-binding protein (BP). In varying compositions they form a membrane spanning structure that can actively change its conformation to facilitate the transport of various compounds through the membrane (Tomii and Kanehisa 1998; Oldham et al. 2008). **Module 79** (4 CLOGs, aAS=0.87) consists of two transmembrane proteins, one ATP-binding protein, and one soluble substrate binding protein comprising an ABC transporter with unclear specificity. **Module 102** (3 CLOGS, aAS = 0.87) groups CLOGs for one substrate-binding proteins as well as two transmembrane proteins for transport of neutral and charged amino acids respectively. These genes are typically found in an operon-like proximity on the genomes together with an ATP-binding protein. **Module 110** (3 CLOGs, aAS=0.62) consists of two transmembrane proteins and one ATP-binding protein forming a putative polyamine transporter; **Module 282** (2 CLOGs, aAS=0.65) consists of a molybdate transporter that is assembled from a fused NBD-TMD protein as well as the substrate-binding protein. Interestingly, in 16 of the 47 strains these two genes are not in close proximity on the genome. **Module 293** (2 CLOGs, aAS=0) consists of the transmembrane and the substrate-binding protein of a carbohydrate transporter. These CLOGs do not form an operon in any strain.

We note that ATP-binding NBD proteins are not always modularized with the corresponding transmembrane and substrate-binding proteins. It is known, that the ATP-binding proteins of different ABC transporters are highly conserved - up to a degree they can functionally substitute each other (Hekstra and Tommassen 1993; Tomii and Kanehisa 1998). The identity of different NBD-proteins can exceed 60% with a BLAST e-value of e-100 and less. The high degree of sequence similarity therefore results in multiple ATP-binding proteins being clustered together in a single CLOG (CLOGs 4775 and 4788), compromising the specific patterns of occurrence.

### Co-occurrences of CLOGs related to metabolic functions

A third class of functional relationships are modules whose constituent CLOGs encode proteins involved in a common metabolic pathway. For example, **module 272** (aAS=0.4) consists of two CLOGs encoding for a phosphoketolase (EC 4.1.2.22) and an acetate kinase (EC 2.7.2.1), respectively. The phosphoketolase catalyzes the reaction of fructose 6-phosphate to erythrose 4-phosphate and acetyl phosphate, the latter is subsequently converted into acetate by the co-occurring acetate kinase. Therefore the module reflects a functional association, although both genes are not in close genomic proximity in 32 of 53 strains, including *Synechocystis* sp. PCC 6803. **Module 203** (aAS=0.9) consists of two CLOGs that code for enzymes of the trehalose synthesis pathway, namely maltooligosyl trehalose synthase (EC 5.4.99.15) and maltooligosyl trehalose hydrolase (EC 3.2.1.141) (Higo et al. 2006). **Module 65** (5 CLOGs, aAS=0.31) is associated with the synthesis of hopanoids, which organize the lipid fraction of cell membranes (Kannenberg and Poralla 1999). The module consists of CLOGs whose genes code for the hopanoid-associated sugar epimerase (HpnA), hopene-associated glycosyltransferase (HpnB), squalene-hopene cyclase (HpnF), hopanoid biosynthesis associated radical SAM protein (HpnH) and a not further specified phosphorylase. All strains participating in this module also harbor at least one copy of the squalene synthase (EC 2.5.1.21), which exists in two variants and is therefore split into the CLOGs 10423 and 10424. **Module 224** (2 CLOGs, aAS=0.91) consists of two CLOGS encoding an aldehyde decarbonylase (Khara et al. 2013) and an acyl-ACP reductase (Schirmer et al. 2010), respectively. The strict co-occurrence and operon-like structure of both genes has already been described in the context of cyanobacterial alkane biosynthesis (Klähn et al. 2014). **Module 303** (aAS=0.9) consists of two CLOGs, acyl-CoA dehydrogenase (EC 1.3.8.7) and 3-hydroxyacyl-CoA dehydrogenase (EC 1.1.1.35) – both integral components of the degradation of fatty acids and branched-chain amino acid. Interestingly, in all cyanobacterial strains, genes of these enzymes form an operon-like structure around a gene which is either part of CLOG 19506 (no clear annotation) or CLOG 9342 (acetyl-CoA acetyltransferase, EC 2.3.1.16). The latter is part of the fatty acids degradation pathway, indicating a similar function of the genes in CLOG 16506. **Module 338** (2 CLOGS, aAS=0.35) consists of the enzymes glucosylglycerol-phosphate synthase (EC 2.4.1.213) (Marin et al. 1998) and glycerol-3-phosphate dehydrogenase (EC 1.1.5.3), which are involved in the synthesis pathway of osmoprotective compound glucosylglycerol (Hagemann and Erdmann 1994). In seven of the 23 strains harboring both CLOGs, the corresponding genes are found in operon-like proximity. Recently, a gene with glycosylglycerol hydrolase activity was identified in Synechosystis sp. PCC 6803 (Savakis et al. 2016). The gene is not part of the module, as the respective CLOG is annotated in only 13 of the strains considered here, and hence does not strictly co-occur.

### Co-occurrences of CLOGs related to specific cellular functions

The final class of modules combines CLOGs related to specific cellular process. For example, **Module 52** (6 CLOGS, aAS=0.34) consists of CLOGs encoding molybdenum cofactor biosynthesis protein A and C, molybdopterin biosynthesis MoeA and MoeE protein as well as a nitrate reductase and a nitrate reductase associated protein. The co-occurrence can be explained by the co-factor molybdopterin providing molybdenum to the reaction center of the nitrate reductase (Woodard et al. 1990). **Module 24** (8 CLOGs, aAS=0.65) consists of 8 CLOGs related to the assembly of gas vesicles proteins. Gas vesicles allow cyanobacteria a controlled lateral movement in liquid medium. The module also contains CLOGs coding for two ATPases with unknown function that might be involved in vesicle formation or pumping processes. The genes of this module are found in close genomic proximity in 10 of the 16 participating genomes.

### Modules provide novel hypotheses for gene function

Of particular interest are modules that combine CLOGs with known function and CLOGs encoding for unknown or putative regulatory proteins. Modules indicating such co-occurrences provide hypotheses about the possible functional role of genes with unknown function and might provide additional insight into regulation of cellular processes. For example **module 111** (3 CLOGs, aAS=0.07) consists of glutamate-5-semialdehyde dehydrogenase (EC 1.2.1.41) involved in the synthesis of essential amino acid L-proline as well as two CLOGs with likely regulatory functions, a pentapeptide repeat protein and a serine/threonine protein kinase. **Module 9** (21 CLOGs, aAS=0.87) contains multiple CLOGs associated with the fixation of inorganic nitrogen, as well as five likely regulatory genes. In *Nostoc* sp. PCC 7120 these are asr1405 (hypothetical protein), all1432 (UBA/THIF-type binding protein, probable hesA), asl1434 (rop-like domain protein), all2512 (probable transcriptional regulator PatB), and asr2523 (TPR domain protein). The putative regulatory genes are located almost always in close genomic proximity to the other genes of module, suggesting a role of these genes in the process of nitrogen fixation.

Other modules involve only CLOGs of unknown function and therefore lack a straightforward functional interpretation. In this case, the shared traits of the strains in which the CLOGs co-occur may provide additional information. For example, **module 3** (aAS=0.24) is comprised of 36 CLOGs mostly annotated as hypothetical or kinase proteins with 7 signaling related proteins, two heterocyst differentiation proteins, four membrane transporter related proteins, and two segregation proteins. Genes of module 3 can, with a few exceptions, only be found in filamentous cyanobacterial strains, indicating a role of these genes in filamentous growth. Likewise, **module 4** (aAS=0.11) combines 30 CLOGs that are solely associated to filamentous cyanobacteria capable of differentiating to heterocysts. The majority of CLOGs in the module lack a specific annotation with only few exceptions, including cytochrome b6f subunit PetM or the heterocyst differentiation protein PatN. Multiple other modules reveal interesting associations of CLOGs, such as CRISPR-related proteins in module 54, 93, and 97, possible chemotaxis genes in module 62, phosphonate lyase related proteins in module 71, and six transposases in module 50.

To facilitate further analysis, we therefore provide the CLOG Similarity Viewer. The viewer includes the complete dataset of co-occurrence and allows the exploration of the co-occurrence neighborhood for any (cyanobacterial) gene of interest. See METHODS for details. Overall, we conclude that the analysis of co-occurrences using a network perspective establishes a suitable tool to generate novel hypotheses about putative functional roles of genes.

## DISCUSSION

Decreasing costs for nucleotide sequencing results in an increased availability of complete genome sequences from all domains of life. The increased coverage of individual phyla offers unprecedented possibilities to investigate and eventually understand the manifold adaptations and functional relationships that determine the genetic content of individual species. In this work, we used comparative genome analysis to investigate the cyanobacterial pan-genome based on 77 strains whose complete genome sequence is available. Our focus was the co-occurrence of likely ortholog genes, denoted as CLOGs – motivated by the hypothesis that co-occurrence is indicative of a functional relationship between CLOGs.

Cyanobacteria form a distinct phylogenetic clade and show enormous diversity in their growth environment (with respect to temperature, salt concentration, humidity), cell shape (single celled, filamentous), and metabolic capability (hydrogen production, diazotrophy). Yet cynaobacteria also (almost) all share a basic metabolic lifestyle, namely autotrophic growth using oxygenic photosynthesis as a primary source of energy and redox potential. We argue that this similar-yet-diverse nature of cyanobacterial growth represents an ideal test case to evaluate gene co-occurrences with respect to putative functional relationships.

To investigate co-occurrences of CLOGs, we introduce a novel approach based on a network perspective that allows us to identify modules of co-occurring CLOGs. Our results demonstrate that modules of co-occurring genes are indeed often indicative of functional relationships. Straightforward examples include known operon-like structures and enzymes that catalyze sequential steps in metabolic pathways. Modules of co-occurring genes, however, can also be associated with specific cellular functions and consist of different genes pertaining to that function. Examples discussed here were the assembly of gas vesicles proteins and the biosynthesis of molybdopterin, among several others. Overall, the individual modules exhibit high functional coherence and provide valuable insight into the functional neighborhood of genes. In this respect, co-occurrence supplements (conserved) co-localization as an indicator of a functional relationship.

We hypothesize that the high functional coherence observed for the extracted modules is also a consequence of the restriction to the similar-yet-diverse cyanobacterial phylum. Related earlier studies either focused on a set of very closely related organisms, such as a set of 8 commensal and 21 pathogenic E. coli strains (Vieira et al. 2011). Or, vice versa, considered global gene-gene co-occurrence in vast number of bacterial species (Kim and Price, 2011). In the former case, the set of organisms typically lacks the necessary diversity to associate genetic content to particular cellular functions and its diversity does not manifest itself in the presence or absence of individual genes. In the latter case, involving a vast number of unrelated bacterial species, we conjecture that possible functional relationships are obscured by the diversity of lifestyles and metabolic functions within the set of considered species. A comparison with the results of a recent analysis involving ∼ 600 bacterial species indeed suggests that the modules identified here are more specific and typically relate to aspects of cyanobacterial functioning and growth, such as nitrogen fixation or formation of gas vesicles – whereas co-occurrence using global analysis primarily revealed examples related to co-occurrence of enzymes related to few basic metabolic pathways (Kim and Price 2011).

Based on our results, we expect that a network analysis of modules of co-occurring CLOGs will become a valuable tool to understand the functional relationships between CLOGs. In this respect, of particular interest are modules that consist of CLOGs with specific annotation, as well as genes with unknown function. Such modules provide novel hypotheses for putative gene functions. To facilitate further analysis based on the results presented here, we therefore provide a computational toolbox, the CLOG Similarity Viewer, which allows identification and exploration of gene-gene co-occurrences beyond the examples discussed in the main text. The toolbox is available for MATLAB (The MathWorks, Inc) as well as a stand-alone application for Mac, Linux, and Windows [http://sourceforge.net/p/similarityviewer/].

## METHODS

### Acquisition of Genomic Data

We searched the NCBI Genome database (http://www.ncbi.nlm.nih.gov/genome/) for cyanobacterial entries and selected all fully sequenced and assembled strains. We further included all associated plasmid sequences, as well as the recently annotated *Escherichia coli* O111:H (denoted as E. coli in the following. In total 78 chromosomes and 136 plasmids were sourced from the NCBI Genome database (January 17. 2015) as listed in Supplement Table 3. An overview of general information of all strains is provided in Supplement Table 4. A phylogenetic tree (Supplement Figure S1) was constructed by extracting the 16S ribosomal RNA sequences of all genomes. Pair wise distances were calculated using the distance model by Jukes and Cantor (Jukes and Cantor 1969) and the BLOSUM62 scoring matrix. The tree was constructed with the “seqlinkage” function by MATLAB using the standard parameter. As expected, the only non-photosynthetic organism Escherichia coli appeared as outgroup.

### Cluster of Likely Orthologous Genes (CLOGs)

Identification of orthologous genes was done as previously described in (Beck et al. 2012). Following an all-against-all BLAST comparison, the bidirectional hit rate between all gene pairs a and b was computed as

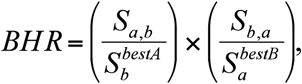

where is the blastp score of *a* versus *b* and 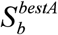 is the best score of *b* against any gene in strain A (including *a*). BHR is one for all mutually best hits and lower otherwise. All gene pairs with a BHR above 0.95 are grouped together. To avoid weakly connected groups, genes in each group were clustered according to their mutual BLAST score using the UPGMA (unweighted pair group method with arithmetic mean) and a cut-off of 20, Hereinafter these clusters are referred to as CLOGs (Cluster of Likely Orthlogous Genes). Accordingly, genes in one cluster are thought to be orthologous and a similar function is assumed. The method was previously evaluated and compared against other available toolboxes and yields similar results (Beck et al. 2012).

### Similarity of CLOGs and modularization

The pair-wise co-occurrence of CLOGs was evaluated in a two-step process. For all 1.7 × 10^9^ pairs of CLOGs, a right-sided Fisher’s exact test was calculated. P-values were corrected for multiple testing using the method by Benjamini and Yekutieli (Benjamini and Yekutieli 2001) with an excepted false discovery rate of 0.01. The critical p-value was below 1.43e-6. For all significantly correlated pairs of CLOG i and j, their similarity was computed as:

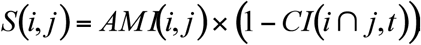

Here, *AMI*(*i.j*) denotes the adjusted mutual information between the sequences of species participating in the CLOGs *i* and *j*, a variant of the mutual information adjusted for lopsided frequencies (Vinh et al. 2010). The AMI ranges between zero for uncorrelated and one for fully correlated pairs. Consistency index *CI*(*I ⋂ j,t*)measures the consistency of the 16S rRNA phylogenetic tree *t* - shown in Supplement Figure S1 - to the set of strains participating in both CLOGs *i* and *j* (Kluge and Farris 1969). Escherichia coli and Cyanobacterium UCYN-A were not considered when calculating the CI.

Anti-correlation between two CLOGs was quantified with the same method but using a left-sided Fisher’s exact test. Correction for multiple testing yielded a critical p-value of 4.42*10^-7^. The consistency index was not computed for anti-correlated CLOGs and therefore set to zero. The adjusted mutual information remains a positive value with plus one for fully anti-correlated pairs. CLOGs were grouped into modules by constructing a graph with nodes representing the CLOGs and weighted edges between positively correlated CLOGs with an AMI of at least 0.65. The heuristic and parameter-free community detection algorithm by Blondel et al. was used to identify modules (Blondel et al. 2008).

### Computation of genomic adjacency

The genomic proximity of genes in one module was calculated for each strain by sorting these genes according to their position on the chromosome or plasmids. Two genes are defined to be in close proximity if less than ten annotated open reading frames separate their loci. By definition, genes on different chromosomes/plasmids are not in close proximity. The adjacency score is then defined as:

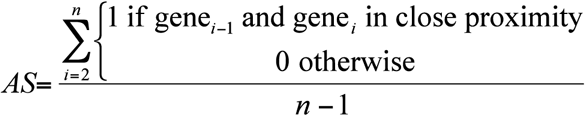

where n is the number of genes within one module. The adjacency score ranges between zero if all genes are spread across the genome and one if they all are in close proximity. The *average AS* (aAS) of a module is then computed as the mean of the AS of all strains with a total of at least two genes in all CLOGs comprising this module.

## DATA ACCESS

Supplement Text 1: PDF file with supplementary methods and figures, tutorial of the Similarity Viewer, examination of the anti-correlation of CLOGs, and a more detailed analysis of the genomic adjacency.

Supplement Table 1: Table of correlated CLOGs sorted by p-value of Fisher’s exact test

Supplement Table 2: Table of all CLOGs sorted by modules.

Supplement Table 3: List of chromosomes and plasmids.

Supplement Table 4: Genomic and environmental information for all strains analyzed in this study.

## ACKNOWLEDGMENTS

**DISCLOSURE DECLARATION** (including any conflicts of interest)

The authors declare no conflict of interest. Primary funding for this study was provided by the research grant CyanoGrowth funded by the German Federal Ministry of Education and Research as part of the e:Bio Innovationswettbewerb Systembiologie [e:Bio systems biology innovation competition] initiative (reference: FKZ 0316192).

